# Impact of Autocalibration Method on Accelerated Echo-Planar Imaging of the Cervical Spinal Cord at 7T

**DOI:** 10.1101/2022.03.08.483513

**Authors:** Alan C. Seifert, Junqian Xu

## Abstract

**Purpose:** The spinal cord contains sensorimotor neural circuits of scientific and clinical interest. However, spinal cord fMRI is significantly more technically demanding than brain fMRI due primarily to its proximity to the lungs. Accelerated EPI at 7T is particularly vulnerable to k-space phase inconsistencies induced by motion or B_0_ fluctuation, during either autocalibration signal (ACS) or timeseries acquisition. For 7T brain fMRI, sensitivity to motion and B_0_ fluctuation can be reduced using a re-ordered segmented EPI ACS based on the fast low-angle excitation echo-planar technique (FLEET). However, respiration-induced B_0_ fluctuations (exceeding 100Hz at C7) are greater, and fewer k-space lines per slice are required, for cervical spinal cord fMRI at 7T, necessitating a separate evaluation of ACS methods.

**Methods:** We compared 24-line single-shot EPI, and 48-line two-shot segmented EPI, two-shot FLEET, and GRE-based ACS acquisition methods, performed under various physiological conditions, in terms of temporal SNR (tSNR) and prevalence of artifacts in GRAPPA-accelerated EPI of the cervical spinal cord at 7T.

**Results:** Segmented EPI and FLEET ACS produce images with nearly identical patterns of severe image artifacts. GRE and single-shot EPI ACS consistently produce images free from significant artifacts, and tSNR is significantly greater for GRE ACS, particularly in lower slices where through-slice dephasing is most severe.

**Conclusion:** GRE and single-shot EPI ACS acquisition methods, which are robust to respiration-induced phase errors between k-space segments, produce images with fewer and less severe artifacts than either FLEET or conventionally segmented EPI for accelerated EPI of the cervical spinal cord at 7T.

## Introduction

The spinal cord contains many neural circuits of scientific and clinical interest, including sensorimotor reflex arcs, modulation of nociception (1), and central pattern generators that produce rhythmic motor activity related to locomotion (2). However, functional MRI (fMRI) of the spinal cord is significantly more technically demanding than fMRI of the brain. High resolution is required to image blood oxygenation level-dependent (BOLD) signal in the spinal cord, which is approximately 12-15 mm in diameter at its widest point, and whose gray matter structures are 1-3 mm wide. Higher static magnetic field strengths (B_0_) yield improved resolution through increased signal to noise ratio (SNR), and increases BOLD signal overall and the microvascular proportion of BOLD signal in particular in gradient-echo echo-planar imaging (EPI) (3,4).

Single-shot echo-planar imaging (EPI) is the predominant method for spatial encoding and readout in functional (fMRI). EPI achieves high temporal resolution by generating and acquiring an echo train containing a full 2D plane of k-space data after a single excitation. However, the long echo train duration and echo time (TE) leaves EPI vulnerable to geometric distortion and signal loss caused by susceptibility-induced field inhomogeneity, particularly at ultra-high field where susceptibility-induced field inhomogeneities are magnified (5). Distortion and signal loss are particularly severe in the spinal cord at 7T (6), where the vertebral column produces significant static field inhomogeneities in the spinal canal that are not effectively countered by spherical harmonic shim functions. Also because of its anatomical location, signal fluctuations due to respiration, motion, and pulsatile rostro-caudal flow of cerebrospinal fluid (CSF) are particularly strong in the spinal cord (7,8). Respiration-induced B_0_ fluctuations (9) can exceed 100Hz at C7 (10). The effects of motion associated with swallowing are also far more detrimental to image quality in the spinal cord.

Geometric distortion and signal loss can be mitigated by reducing the length of the EPI echo train by parallel imaging. For this reason, GRAPPA-accelerated (11) EPI is a mainstay in both brain and spinal cord BOLD fMRI at ultra-high field. Acquisition of autocalibration signal (ACS) data that is uncorrupted by motion or B_0_ fluctuations is crucial for maximizing temporal signal to noise ratio (tSNR) and minimizing the effects of artifacts in timeseries fMRI data (12). However, image contrast in the autocalibration signal (ACS) dataset need not be precisely matched to image contrast in the accelerated timeseries dataset (13), leaving some valuable flexibility in the choice of ACS acquisition method that can be leveraged to address the unique challenges presented by temporal B_0_ fluctuations in GRAPPA-accelerated spinal cord fMRI.

For 7T brain fMRI, ACS data is often acquired using either a single-shot or segmented multi-shot EPI acquisition. Single-shot EPI is robust to motion or temporal fluctuation of B_0_ because it acquires ACS data for a given slice as a single ‘shapshot’ in time. Multi-shot EPI acquires a larger ACS dataset, reducing the effects of noise on GRAPPA kernel calculation and increasing the SNR of the accelerated EPI dataset, but is susceptible to k-space phase inconsistencies resulting from differences in respiratory phase or motion between the multiple shots. Polimeni et al. (14) demonstrated reduced sensitivity to motion and B_0_ fluctuation using a re-ordered segmented EPI ACS based on the fast low-angle excitation echo-planar technique (FLEET) (15). In FLEET, all shots in a multi-shot EPI ACS acquisition are completed in quick succession for a given slice before proceeding to the next slice, increasing robustness to potential motion, but reducing signal amplitude due to relatively less recovery of longitudinal magnetization between shots. Nevertheless, FLEET was shown to outperform segmented multi-shot EPI, single-shot EPI, and even non-EPI gradient echo (GRE) ACS acquisition (16) methods in terms of tSNR in the brain.

However, in spinal cord fMRI, the relative impacts of respiratory field fluctuations and motion are different, and the number of k-space lines per slice is less, compared to brain fMRI, necessitating a separate evaluation of ACS methods. In this study, we compared single-shot EPI, two-shot segmented EPI, two-shot FLEET, and GRE-based ACS data acquisition methods, performed under various physiological conditions, in terms of tSNR and prevalence of artifacts in GRAPPA-accelerated EPI of the cervical spinal cord at 7T.

## Methods

Imaging was performed on a 7T actively-shielded whole-body scanner (Magnetom, Siemens) using a 22-channel brainstem/cervical spinal cord RF coil (17). All imaging experiments were performed under an IRB-approved protocol and in full compliance with all applicable human subject protection regulations. Shim adjustment was performed during free-breathing. Axial single-shot gradient-echo (GRE)-EPI images of the cervical spinal cord were acquired spanning vertebral levels C4-C7 in four healthy volunteers (ages 26-38, 3 males, 1 female) with a well-tested 7T cervical spinal cord fMRI protocol: 0.75 × 0.75 × 3 mm^3^, FOV = 84 × 84 mm^2^, TR/TE = 1500/20.8 ms, flip angle = 70°, phase encoding AP, GRAPPA R = 2, partial Fourier = 6/8, BW = 1088 Hz/px, echo spacing = 1.04 ms, and one coronal saturation slab anterior to the cord.

Twelve experiments were performed to test four ACS data acquisition methods (24-line single-shot EPI, and 48-line two-shot segmented EPI, two-shot FLEET, and GRE (16), Figure 1) under three physiological conditions during ACS data acquisition (end-expiration breath-hold, free-breathing, and intentional swallowing). FLEET and GRE ACS data acquisition used flip angle = 12°.

**Figure 1:**
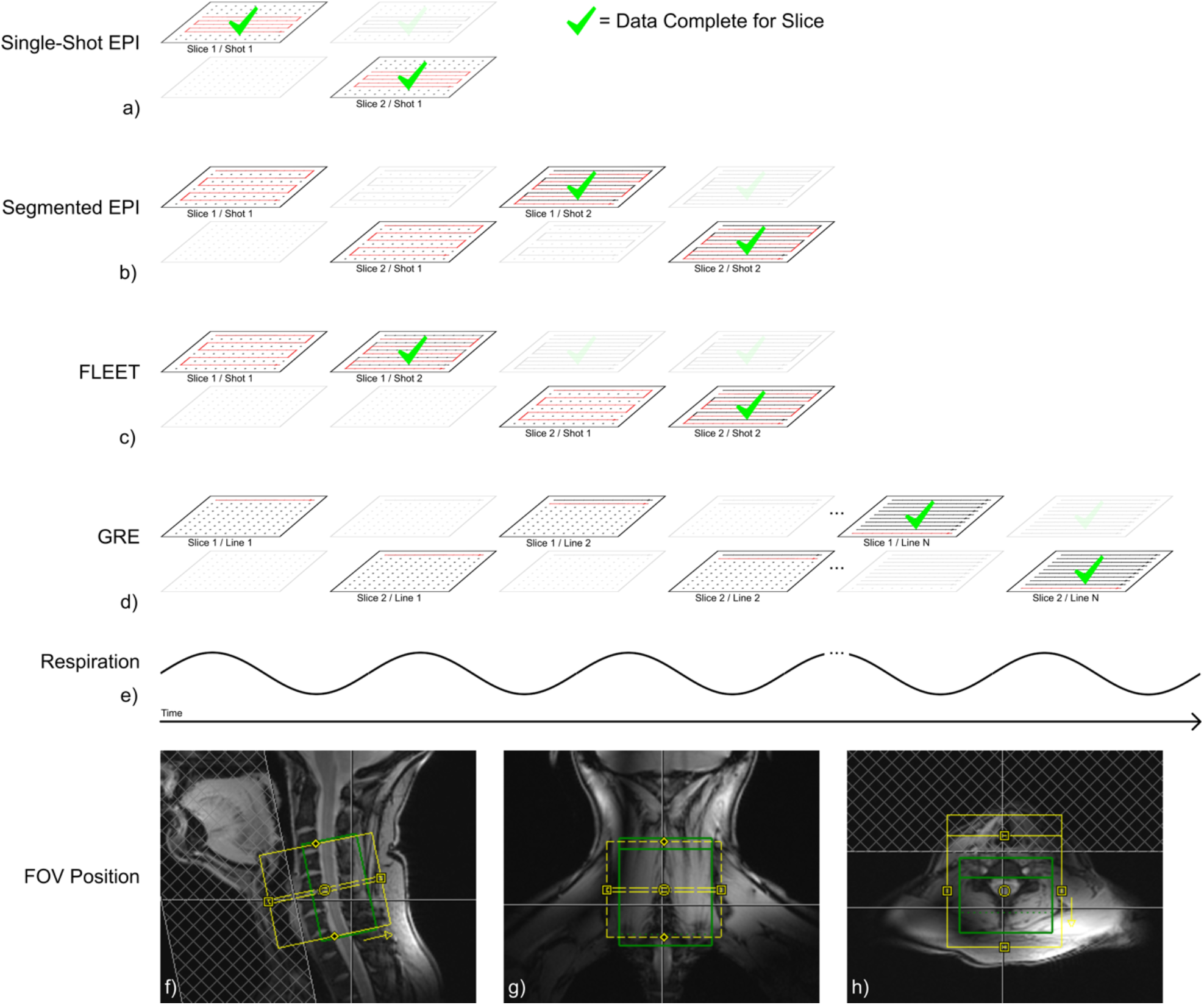
Schematic of k-space data acquisition order of single-shot EPI (a), segmented EPI (b), FLEET (c), and spin-warp GRE (d) autocalibration signal data acquisition. Single-shot EPI acquires each ACS image at a single, discrete phase of the respiratory cycle (e), while segmented EPI and FLEET acquire multiple k-space segments per imaging slice at different phases of the respiratory cycle, leading to phase discontinuity between k-space segments. GRE ACS data also contain phase errors, but these errors are distributed incoherently across a large number of k-space phase-encode lines. The imaging FOV (yellow) and shim adjustment volume (green) are also displayed (f-h).

During imaging data acquisition (i.e., after ACS acquisition), the subject breathed freely and refrained from swallowing for 20 frames (30s), and then intentionally swallowed every 3-5s for the following 20 frames (30s). Off-resonance correction of B_0_ fluctuation between fMRI frames was applied using a vendor provided algorithm integrated in the image reconstruction (18). All image reconstruction was performed on the scanner. Image timeseries were motion-corrected (x- and y-translation) using FSL FLIRT (19). Temporal SNR was calculated voxelwise, and averaged within manually defined masks of the spinal cord for tabulation and statistical analysis. The free breathing and cued swallowing timeseries were treated separately in statistical analyses.

## Results

Temporal mean images and tSNR maps of data acquired using the various ACS methods under the most common scenario of free breathing during both ACS and timeseries data acquisition are displayed in Figure 2. Temporal mean images and tSNR maps under all of the physiological conditions investigated are shown in Figure 3 and plotted in Figure 4. GRE ACS yields the highest average tSNR of 10.08 across all experiments, FLEET yields the second highest of 9.16, single-shot EPI ACS yields the third highest at 7.63, and segmented EPI yields the lowest at 7.08. Pair-wise comparisons of tSNR between ACS acquisition methods and physiological conditions during ACS acquisition are given in Figure 5.

**Figure 2:**
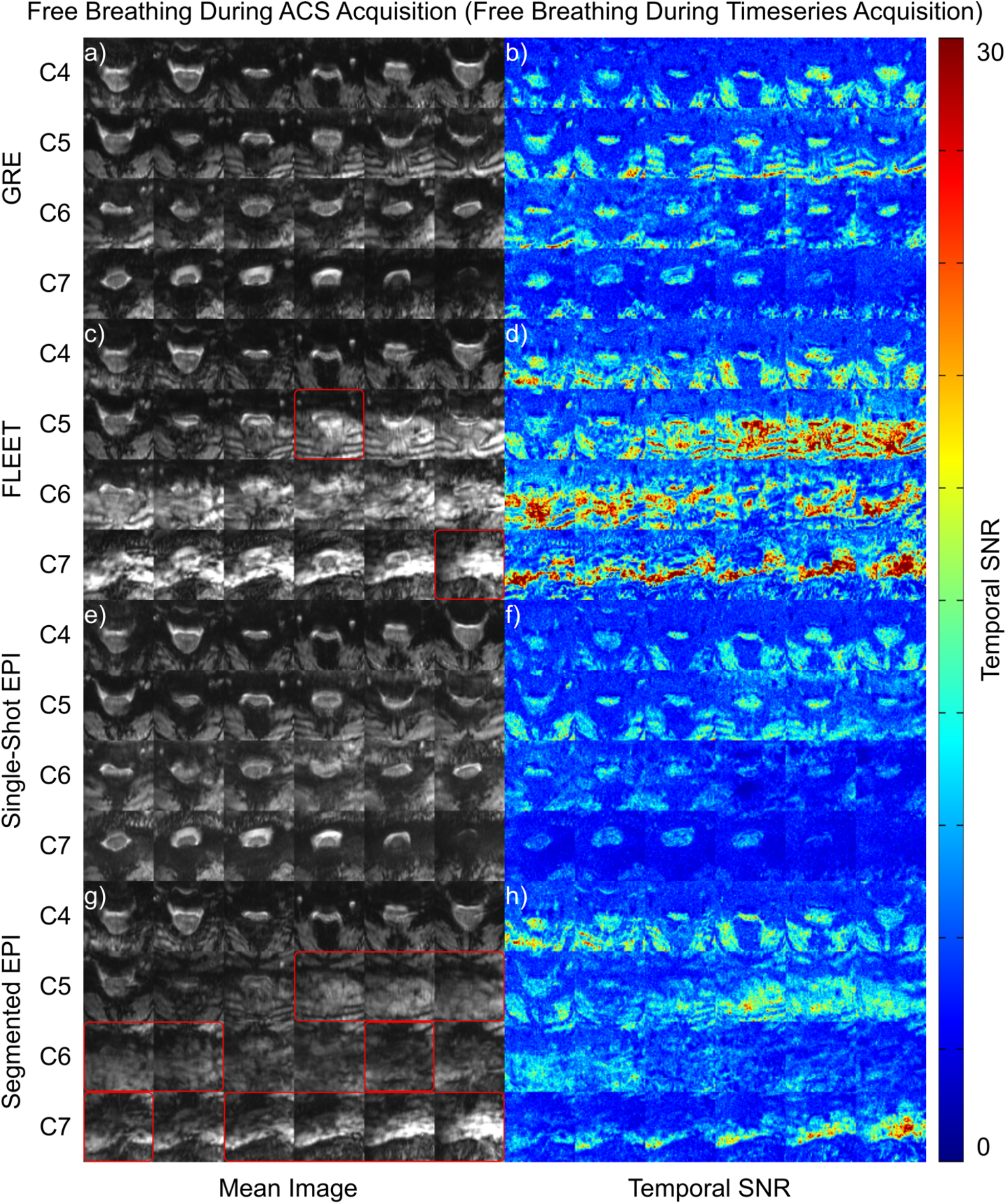
Temporal mean images (a,c,e,g) and temporal signal to noise ratio maps (b,d,f,h) produced by 2D echo-planar imaging acquisitions with four autocalibration signal (ACS) acquisition methods in a single representative subject. Both ACS and timecourse acquisitions are performed under free breathing with no swallowing in this montage, representative of the most common scenario in in vivo scans. GRE and FLEET ACS produce images with high temporal signal to noise ratio, while GRE and single-shot EPI ACS methods produce images largely free of artifacts. Red boxes indicate slices that are unusable due to severe artifacts; these are excluded from tSNR plots and statistical tests.

**Figure 3:**
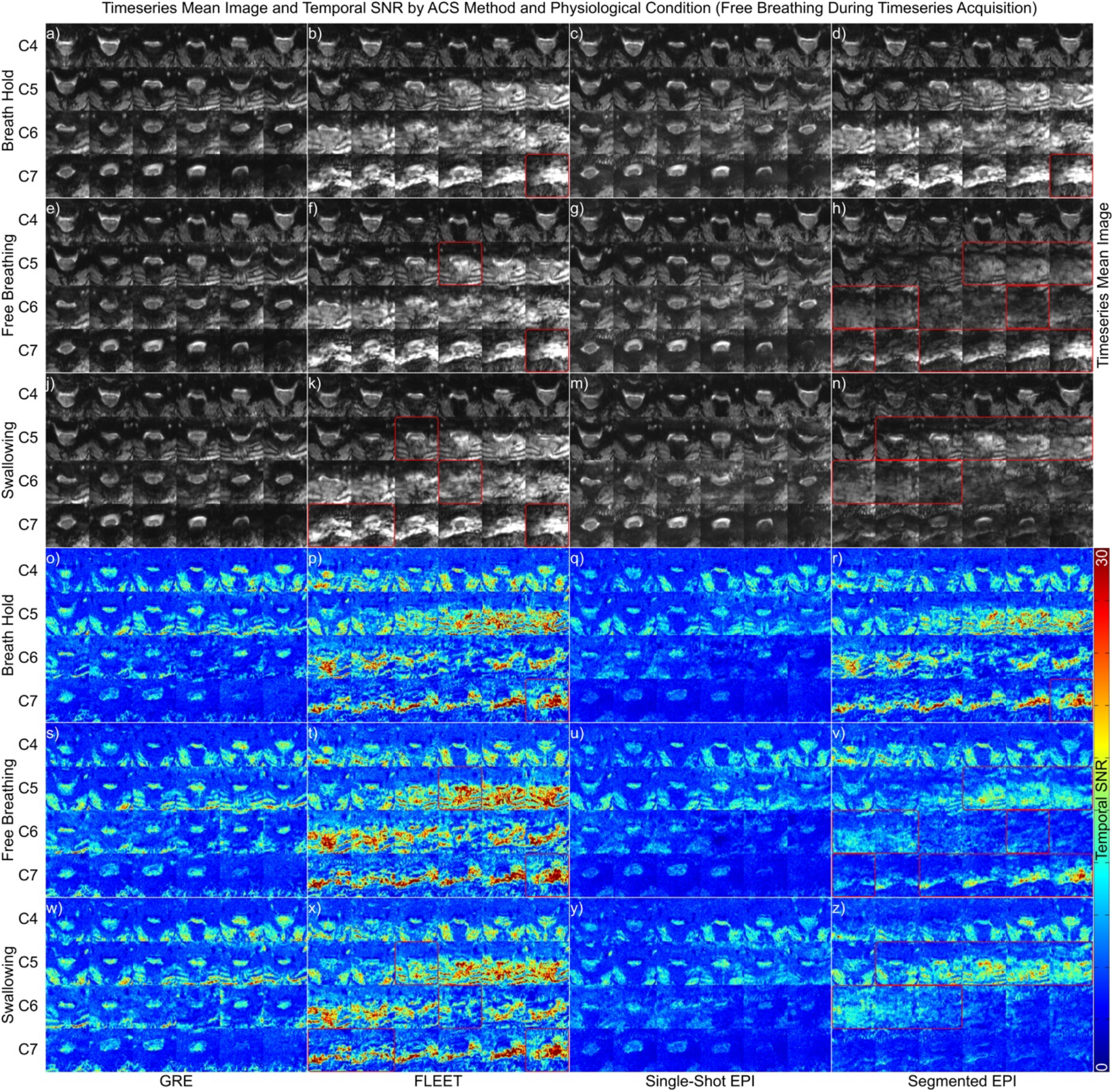
Temporal mean images (a-n) and temporal signal to noise ratio maps (o-z) produced by 2D echo-planar imaging acquisitions with four autocalibration signal (ACS) acquisition methods in a single representative subject. ACS acquisition is performed under three physiological conditions: breath hold, free breathing with no swallowing, and free breathing with a single swallow. Timecourse acquisitions were performed under free breathing with no swallowing, and with cued swallowing, though only free breathing images and maps are displayed here. Red borders indicate that a given slice is so corrupted by artifacts as to be rendered unusable.

**Figure 4:**
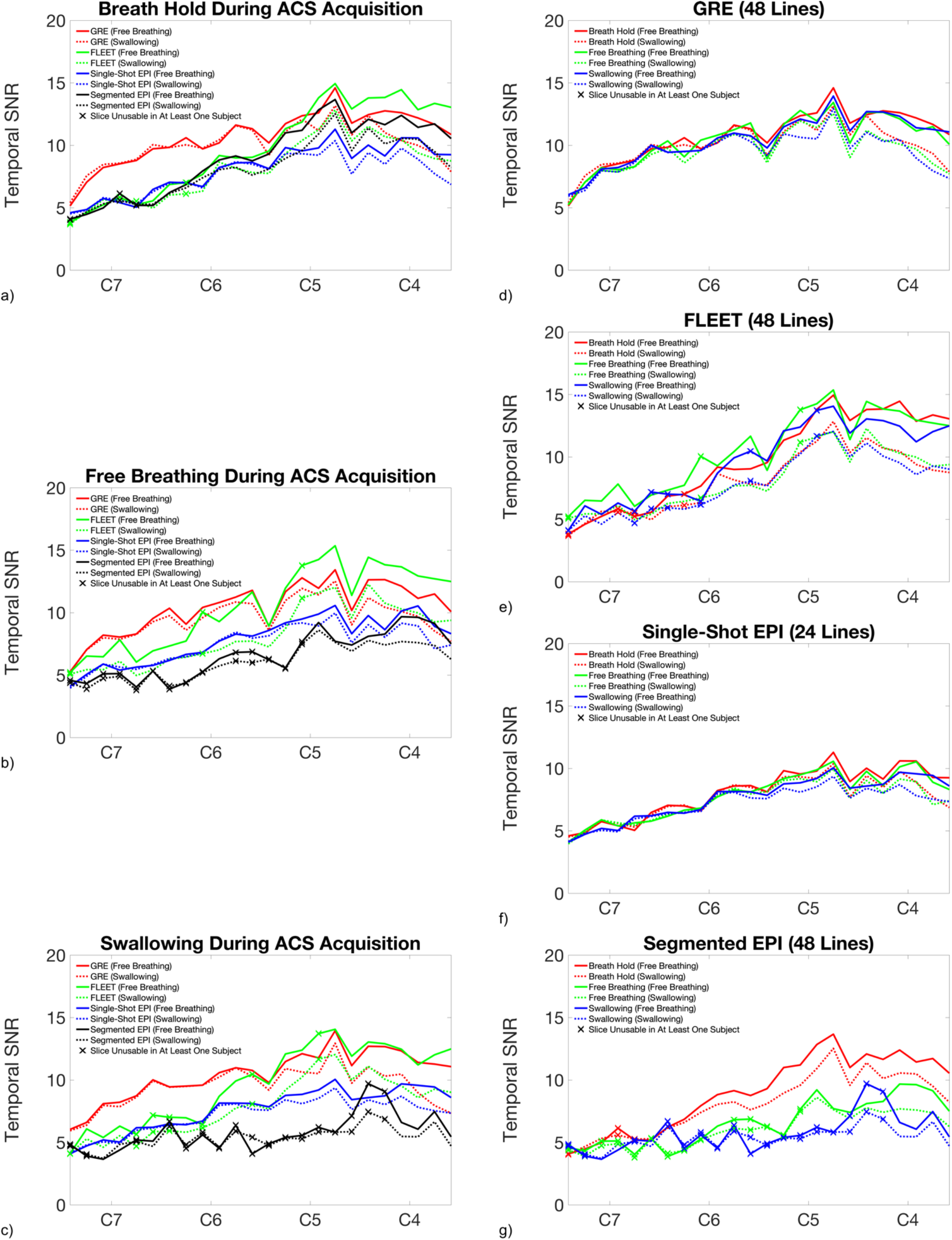
Slicewise temporal signal to noise ratio averaged within the spinal cord across all subjects for each of the three physiological conditions during autocalibration signal (ACS) acquisition (a-c) and four ACS acquisition methods (d-g). In (a-c), data are plotted in different colors for the four ACS acquisition methods. In (d-g), data are plotted in different colors for three physiological conditions during ACS acquisition (breath hold, free breathing with no swallowing, and free breathing with a single swallow). In all frames, solid and dashed lines correspond to the two physiological conditions during timeseries acquisition: free breathing with no swallowing, and free breathing with swallowing, respectively. FLEET and GRE ACS acquisition methods consistently produce high temporal signal to noise ratio in more superior slices, while GRE outperforms the other ACS acquisition methods in more inferior slices. In breath-held ACS acquisitions, the three EPI-based methods (single-shot, segmented, and FLEET) perform similarly, but vary in their robustness to breathing and swallowing: segmented EPI and, to a lesser extent, FLEET ACS acquisition methods are affected by breathing and swallowing during ACS acquisition, while single-shot EPI (as well as GRE) are relatively robust to these perturbations.

**Figure 5:**
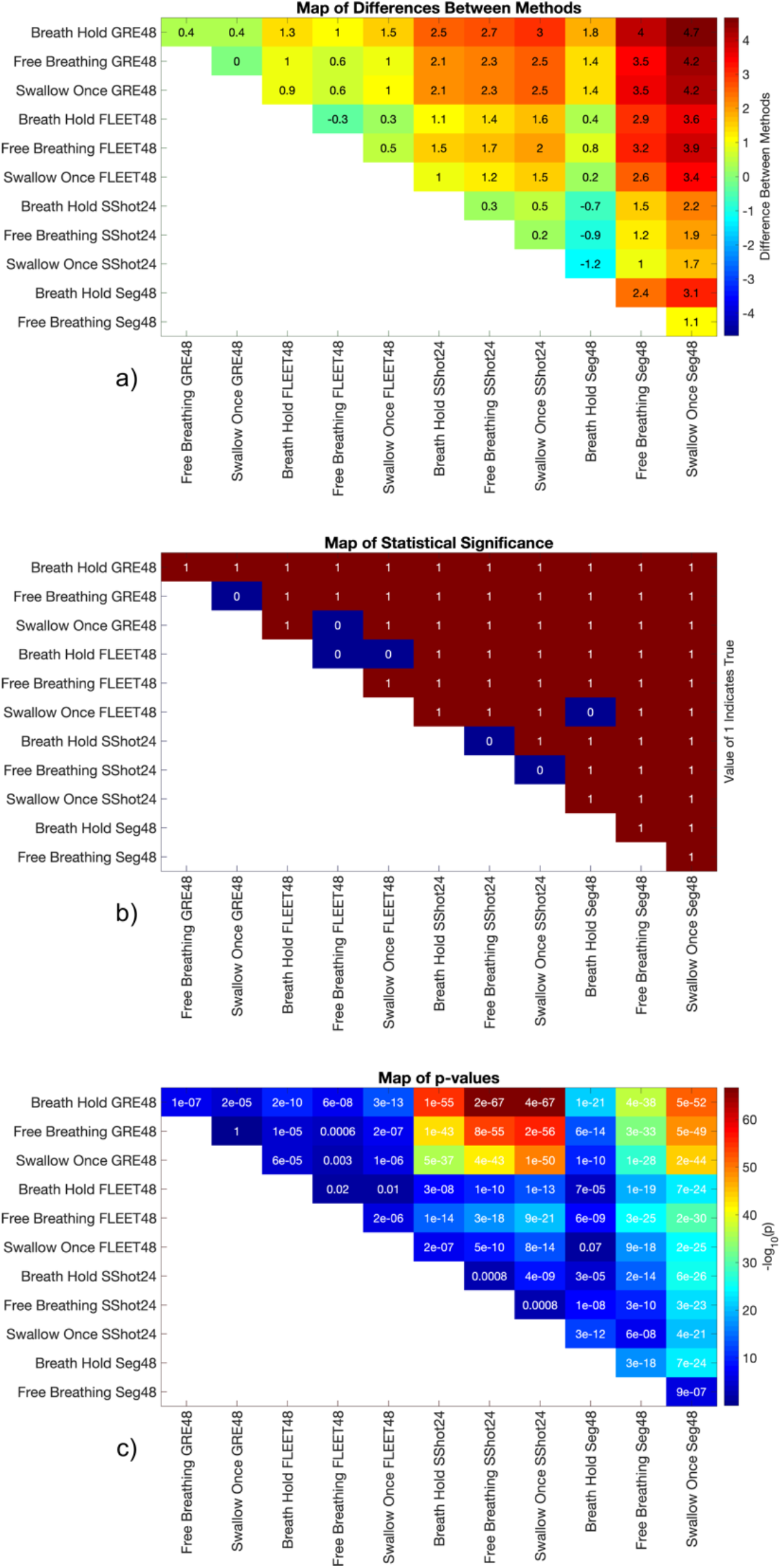
Pair-wise comparisons of temporal signal to noise ratio (tSNR) between autocalibration signal (ACS) acquisition methods and physiological conditions during ACS acquisition. (a) Differences in tSNR between the methods and conditions indicated by the row and column labels (e.g., tSNR in breath hold GRE48 is 0.4 greater than tSNR in free breathing GRE48). (b) Statistical significance and (c) p-values of paired two-sample t-tests between each pair of methods and conditions. A Bonferroni-corrected p-value threshold of 7.58×10^−4^ was used as the criterion for statistical significance. If a slice in a given method was unusable, then the corresponding slice in all other methods was excluded in comparisons; thus, these results compare tSNR only, and do not fully represent differences in image artifacts between methods. Temporal SNR measurements under free breathing with no swallowing and free breathing with swallowing are treated as separate observations.

GRE and single-shot EPI ACS consistently produce images free from significant artifacts. Of these two, GRE ACS produces higher tSNR, particularly in more inferior slices where respiration-induced dynamic field inhomogeneity is more severe.

FLEET and segmented EPI ACS resulted in a greater prevalence of severe artifacts in lower slices that obliterate the anatomical detail of the spinal cord. These artifacts vary among physiological conditions during ACS acquisition using segmented EPI, but are consistent using FLEET. For FLEET ACS, 17 of 288 total slices were unusable due to severe artifacts overlaid on the spinal cord; for segmented EPI ACS, 45 of 288 total slices were unusable.

Cued intentional swallowing during timeseries acquisition decreased tSNR for all ACS methods.

## Discussion

In the cervical spinal cord, especially in lower vertebral levels closer to the lungs, respiration-induced field shifts are a far more prominent source of image artifacts than motion. Respiration generally produces B_0_ fluctuations exceeding 100Hz at 7T (10) continuously throughout acquisition of ACS data, and these fluctuations occur on a similar time-scale to TR. Therefore, unlike in whole brain fMRI (14), where isolated instances of head motion are the primary concern, severely disruptive phase errors between segments are approximately as likely in FLEET ACS as in conventionally segmented EPI ACS for cervical spinal cord fMRI. Indeed, these two methods exhibit nearly identical patterns of image artifacts (Figures 2-3). The detrimental effect of these phase errors is illustrated by the fact that image artifacts are more visually apparent in images reconstructed using two 24-line segments of ACS data (segmented EPI and FLEET) than a single 24-line segment (single-shot EPI). In other words, phase inconsistency is not a risk whose occurrence is minimized by acquiring all shots for a slice as close together in time as possible; phase inconsistency during multi-shot acquisition in the spinal cord is a near-certainty, and is approximately as likely to occur in FLEET as in conventionally segmented EPI. Although phase error correction in segmented EPI ACS lines is a potential research area (20), such advanced image reconstruction methods are beyond the scope of this manuscript.

GRE and single-shot EPI ACS acquisition methods, on the other hand, appear largely free of image artifacts. This suggests that incoherence of phase errors across acquisition of many spin-warp GRE ACS k-space lines (with minimum TE) produces far less detrimental effects on images than phase errors of a similar magnitude but distributed across a smaller number of shots (two, in the present case) as in segmented EPI and FLEET ACS. Although the calculated tSNR in segmented EPI and FLEET ACS exceeds that of single-shot EPI ACS, much of the image signal in segmented EPI and FLEET arises from artifacts rather than the spinal cord, confounding the tSNR measurements.

GRE ACS produces images qualitatively similar to those reconstructed using single-shot EPI ACS, with the least sensitivity to intentional swallowing during timeseries data acquisition, and with the highest tSNR, particularly in lower slices where through-slice B_0_ gradients are the strongest.

It is generally understood to be the best practice to acquire ACS data using the same k-space readout method as the main imaging data, especially in cases where susceptibility-induced distortion is severe, such as the spinal cord. Doing so ensures that patterns of spatial distortion are matched (21,22). However, it has been shown that GRE-based acquisition of ACS data for EPI results in improved tSNR in the brain by dramatically reducing sensitivity to phase inconsistency (or, rather, rendering it incoherent) in the ACS data primarily due to bulk head motion (16), which resulted in its adoption in the Human Connectome Project 7T brain fMRI protocol (23,24). Here we find a similar improvement in the spinal cord at 7T, where susceptibility-induced distortions are particularly severe. The present results suggest that the observed increase in tSNR in the spinal cord using GRE-based ACS acquisition is achieved by the same mechanism as in the brain, and that this improvement outweighs any drawbacks related to a mismatch of spatial distortion patterns between ACS and imaging data.

GRE and single-shot EPI autocalibration signal acquisition methods, which are robust to respiration-induced phase errors between k-space segments, produce images with fewer and less severe artifacts than either FLEET or conventionally segmented EPI for accelerated EPI of the cervical spinal cord at 7T. These findings are expected to generalize to spinal cord fMRI at 3T as well.

## Acknowledgements

This study was supported by National Institutes of Health (NINDS) award K01NS105160 (ACS) and Department of Defense MSRP grant W81XWH-17-1-0204 (JX). Data in this study was acquired using the University of Minnesota Center for Magnetic Resonance Research (CMRR) Multiband EPI pulse sequence software package. A version of this study in a single subject has previously been presented as a conference abstract (25).

## Notes

Disclosures: All authors state that they have no conflicts of interest.

### Competing Interest Statement

The authors have declared no competing interest.

